# Temperate grass allergy season defined by spatio-temporal shifts in airborne pollen communities

**DOI:** 10.1101/410829

**Authors:** Georgina L. Brennan, Caitlin Potter, Natasha de Vere, Gareth W. Griffith, Carsten A. Skjøth, Nicholas J. Osborne, Benedict W. Wheeler, Rachel N. McInnes, Yolanda Clewlow, Adam Barber, Helen M. Hanlon, Matthew Hegarty, Laura Jones, Alexander Kurganskiy, Francis M. Rowney, Charlotte Armitage, Beverley Adams-Groom, Col R. Ford, Geoff M. Petch, The PollerGEN Consortium, Simon Creer

## Abstract

Grass pollen is the world’s most harmful outdoor aeroallergen and sensitivity varies between species. Different species of grass flower at different times, but it is not known how airborne communities of grass pollen change in time and space. Persistence and high mobility of grass pollen could result in increasingly diverse seasonal pollen communities. Conversely, if grass pollen does not persist for an extended time in the air, shifting pollen communities would be predicted throughout the summer months. Here, using targeted high throughput sequencing, we tracked the seasonal progression of airborne Poaceae pollen biodiversity across Britain, throughout the grass allergy season. All grass genera displayed discrete, temporally restricted peaks of pollen incidence which varied with latitude, revealing that the taxonomic composition of grass pollen exposure changes substantially across the allergy season. By developing more refined aeroallergen profiling, we predict that our findings will facilitate the exploration of links between taxon-specific exposure of harmful grass pollen and disease, with concomitant socio-economic benefits.

## Introduction

Allergens carried in airborne pollen are associated with both asthma [1] and allergic rhinitis (hay fever), negatively affecting 400 million people worldwide [2]. Pollen from the grass family (Poaceae) constitutes the most significant outdoor aeroallergen [3, 4], and more people are sensitised to grass pollen than to any other pollen type [5]. However, despite the harmful impact of grass pollen on human health, current observational studies and forecasts categorize grass pollen at the family level [Poaceae; 6, 7] due to difficulties in differentiating species and genera of grass pollen based on morphology [8]. Furthermore, we cannot predict seasonal variation in airborne grass pollen from the phenology of local grasses at ground level, since airborne pollen can be highly mobile [9, 10] and often does not directly correlate to local flowering times [9]. Understanding the taxon-specific phenology of airborne pollen would fill a significant knowledge gap in our understanding of allergen triggers, with associated benefits to healthcare providers, pharmaceutical industries and the public.

Many species within the subfamilies Pooideae, Chloridoideae, and Panicoideae release allergenic pollen into the atmosphere [5], including *Phleum* spp. (e.g. Timothy grasses), *Dactylis* spp. (Cocksfoot grasses), *Lolium* spp. (Ryegrasses), *Trisetum* spp. (Oatgrasses), *Festuca* spp. (Fescues), *Poa* spp. (Meadow-grasses and Bluegrasses), and *Anthoxanthum* spp. (Vernal grasses). However, it is unknown whether particular grass species contribute more to the prevalence of allergic symptoms and related diseases than others [11]. Whilst some grasses have been identified as more allergenic than others *in vitro* (triggering higher levels of Immunoglobulin E (IgE) antibody production), there is a high degree of cross-reactivity between grass species [12]. In addition, the allergen profiles and the degree of sensitisation differ between grass species [12, 13] and the allergenicity of grass pollen varies across seasons [14]. Family-level estimates of grass pollen concentrations cannot therefore be considered a reliable proxy for either the concentration of pollen-derived aeroallergens or pollen-induced public health outcomes.

The identification of biodiversity via the high-throughput analysis of taxonomy marker genes (popularly termed metabarcoding) provides an emerging solution to semi-quantitatively identify complex mixtures of airborne pollen grains [15-18]. Previous metabarcoding studies of airborne pollen have been performed at very limited spatial and temporal scales [e.g. 15, 16]. Recent global DNA barcoding initiatives and co-ordinated regional efforts have now resulted in near complete genetic databases of national native plants, including grasses in Great Britain [19].

Here, using two complementary DNA barcode marker genes (*rbcL* and ITS2), we characterise the spatial and temporal distribution of airborne grass pollen throughout the temperate summer grass pollen season (May-August) across the latitudinal and longitudinal range of Great Britain (S1 Fig). We hypothesise that (i) there will be discrete temporal incidences of pollen from different grasses, linked to Poaceae terrestrial phenology, and (ii) the composition of grass pollen will be homogenous across the UK due to the potential for long distance transport of windborne pollen grains.

## Results and Discussion

Grass pollen occupied distinct temporal windows across the grass allergy season in 2016 (May to August), thereby supporting our hypothesis (i) that species composition of airborne grass pollen will change throughout the grass allergy season (Fig 1, Fig 2). Time, measured as number of days after the first sample was collected, is a good predictor of airborne grass pollen taxon composition using both markers (Fig 1-2; *ITS2*, LR_1, 74_ = 128.8, *P* = 0.001; *rbcL*, LR_1, 71_ = 46.71, *P* = 0.001). We found that month (coded as a factor in the models) improves our ability to predict taxonomic composition across the pollen season (Fig 1-2; *ITS2*, LR_1, 70_ = 319.7, *P* = 0.001; *rbcL*, LR_1, 67_ = 217.25, *P* = 0.001). In addition, community-level ordination reveals that the community as a whole changed across the allergy season (S2 Fig).

**Fig 1.**
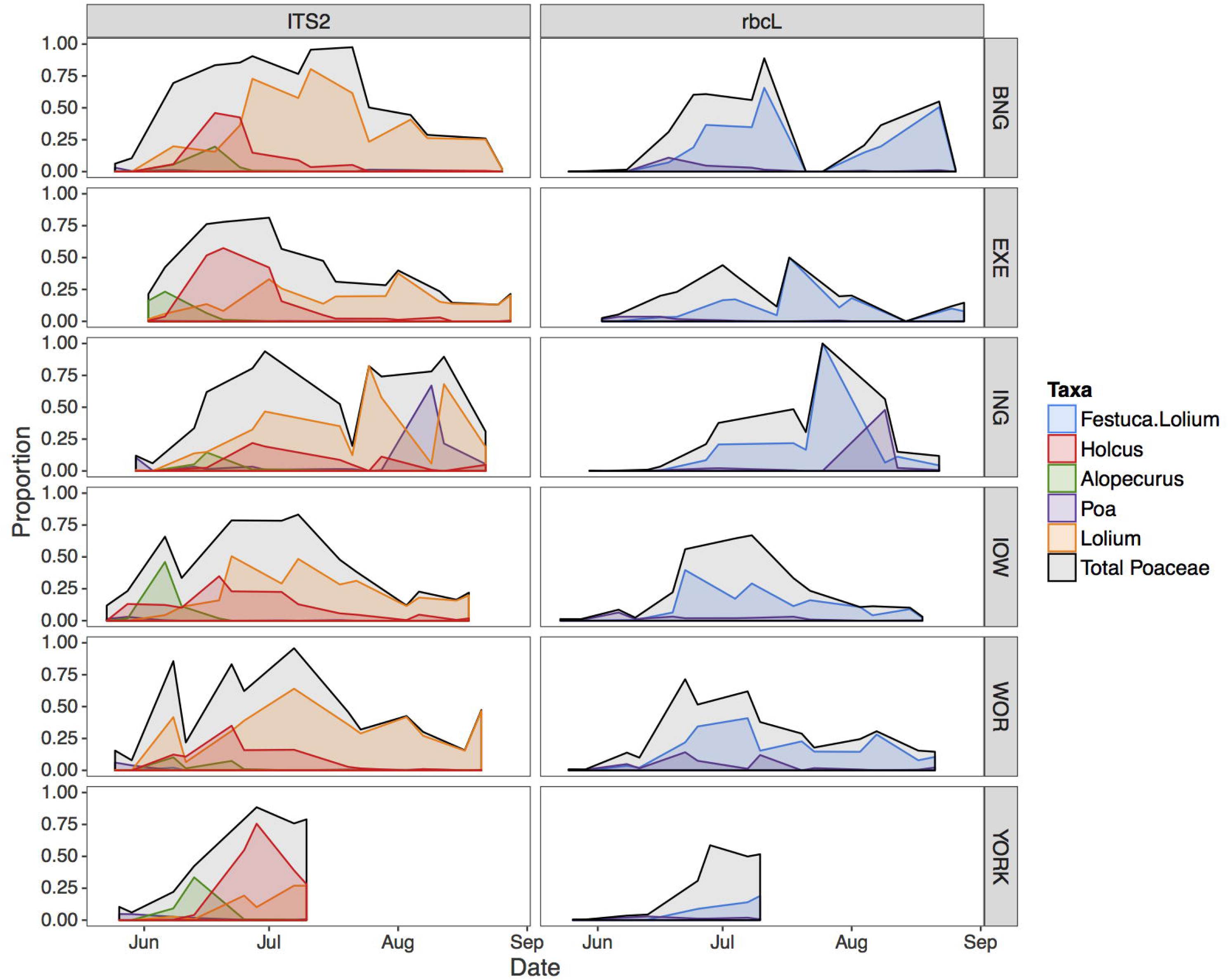
Abundance of the most common airborne grass pollen taxa throughout the grass allergy season. The five most abundant grasses (expressed as proportion of total reads), depicted alongside the total proportion of reads assigned to family Poaceae. Markers used to identify grass pollen are stated in the top panel label. Due to errors in sampling equipment, only 4 weeks of samples were collected at the York sampling site. Sampling sites are indicated in the right panel label abbreviated as follows: BNG = Bangor; EXE = Exeter; ING = Invergowrie; IOW = Isle of Wight; WOR = Worcester; YORK = York. A map of sampling locations can be found in S1 Fig.

**Fig 2.**
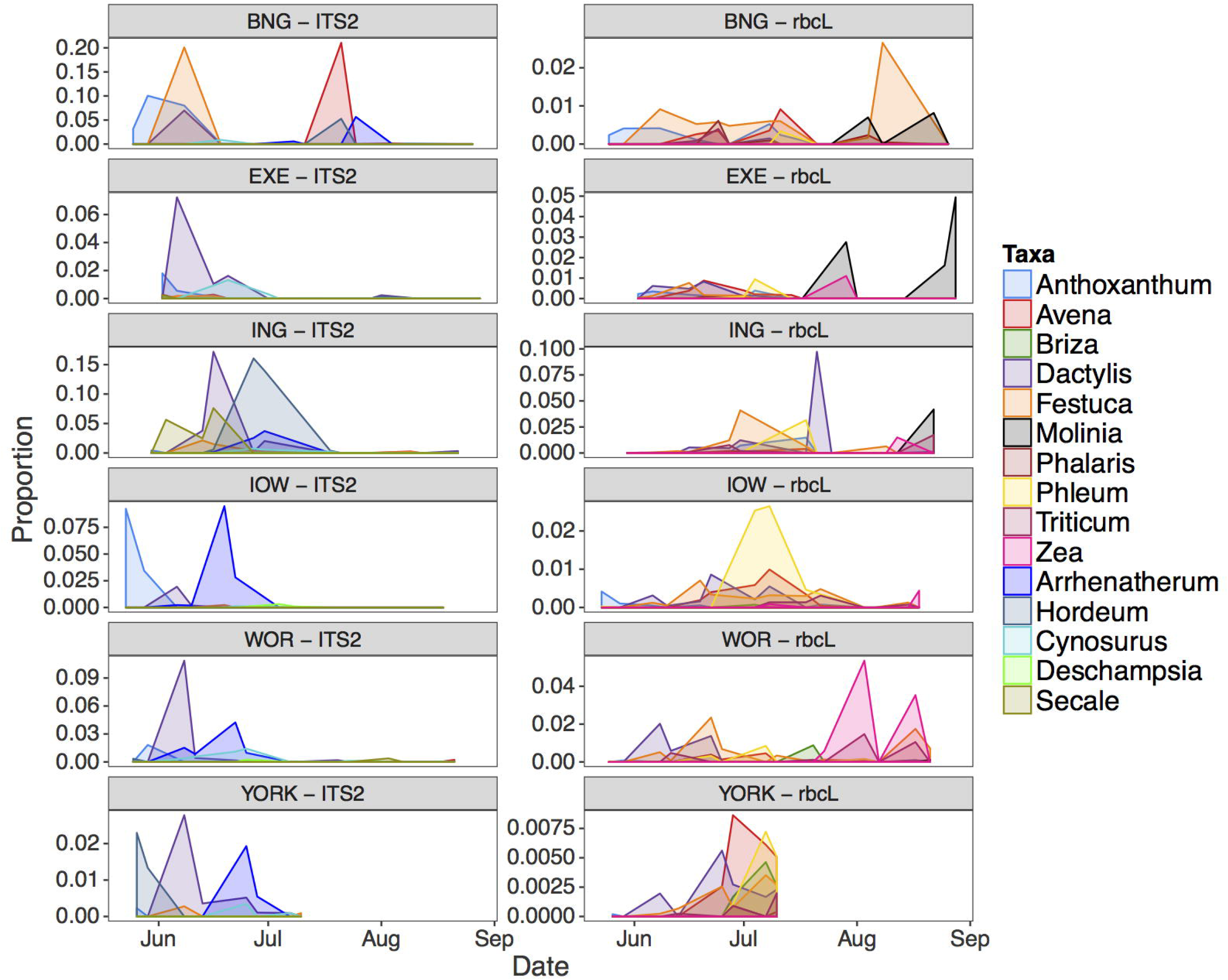
Abundance of airborne grass pollen taxa throughout the grass allergy season. Abundance of rare grasses (expressed as proportion of total reads). Sampling sites are indicated in the top panel, followed by the marker used to identify grass pollen. Due to errors in sampling equipment, only 4 weeks of samples were collected at the York sampling site. Note that the y axes differ between panels. Refer to Fig 1 for site name abbreviations.

Focusing on the more taxonomically specific ITS2 marker dataset, *Alopecurus* and *Holcus* typically dominated the early grass pollen season (Fig 1), which coincides with typical peaks in allergic rhinitis [20], but further research will be required to confirm this association. *Lolium* featured prominently for the majority of the later grass season. The popularity of *Lolium* species as forage crop means that many varieties have been bred with the potential to mature at different times throughout the year [21]. While *Lolium* was the dominant species in airborne grass pollen from July to the end of the sampling period, the total grass pollen concentration declined in August, indicating that the absolute number of *Lolium* pollen grains at this time is low (S3 Fig).

The top five genera contributing to airborne pollen, indicated by the relative abundance of taxonomy marker genes, were *Alopecurus, Festuca, Lolium, Holcus* and *Poa* (Fig 1; S3 Fig). Each of these genera are widespread in the UK and have been shown to provoke IgE-mediated responses in grass-sensitised patients [12], providing candidate species for links with hay fever and asthma exacerbation. Conversely, less prevalent species in the dataset could contribute disproportionately to the allergenic load. Species such as *Phleum pratense* have been identified to be a major source of allergenic pollen [5, 22]. However, we found that *Phleum* made up a very small proportion of metabarcoding reads (Fig 2), corresponding with the results of an earlier phenological study [23]. Most genera, such as *Phleum, Anthoxanthum* and *Dactylis*, show distinct and narrow temporal incidence (Fig 2), and could allow researchers to identify grass species associated with allergenic windows with greater accuracy.

Changes in species composition over time were localised. We found that peaks in abundance of airborne pollen occurred at different times at each location during the summer (Fig 1-2). For example, the relative abundance of airborne grass pollen from the genus *Poa* peaked in mid-June in Worcester and Bangor but 6-8 weeks later in Invergowrie (Fig 1), probably due to latitudinal effects on flowering time [7, 24]. This is supported by a significant interaction between latitude and time of year for both markers (Fig 1-2; *ITS2*, LR_68, 1_= 46.4, *P* = 0.001; *rbcL*,LR_66, 1_= 59.08, *P* = 0.001), and between longitude and time of year for the ITS2 dataset (Fig 1-2; LR_67, 1_= 37.5, *P* = 0.001). Differences in species composition of airborne grass pollen between the six sampling sites is supported by a significant effect of latitude (Fig 1-2; *ITS2*, LR_1, 73_ = 73.2, *P* = 0.001; *rbcL*, LR_1, 70_ = 26.4, *P* = 0.025) and longitude (Fig 1-2; *ITS2*, LR_1, 69_ = 36.5, *P* = 0.005; *rbcL*, LR_1, 69_ = 27.10 *P* = 0.018). These results do not support our hypothesis (ii) that the composition of grass pollen will be homogenous across the UK, and instead suggest taxon-specific effects of regional geography and climate which have been demonstrated for Poaceae pollen as a whole [7].

Observations of first flowering dates from a citizen science project (UKPN; www.naturescalendar.org.uk) and metabarcoding data show similar sequences of seasonal progression (Fig 3). First flowering dates of each genus started almost 3-4 weeks prior to the observation of peaks of grass pollen in the metabarcoding data (Fig 3). Pollen release (anthesis) occurs approximately 2-3 weeks after the production of flowering heads (heading) [25], and this is reflected in the metabarcoding data suggesting that local flowering data are informative for predicting the composition of airborne pollen. Continuing this study over multiple years would allow us to track long-term, phenological changes in airborne pollen communities and improve our ability to forecast the seasonal progression of airborne pollen [26].

**Fig 3.**
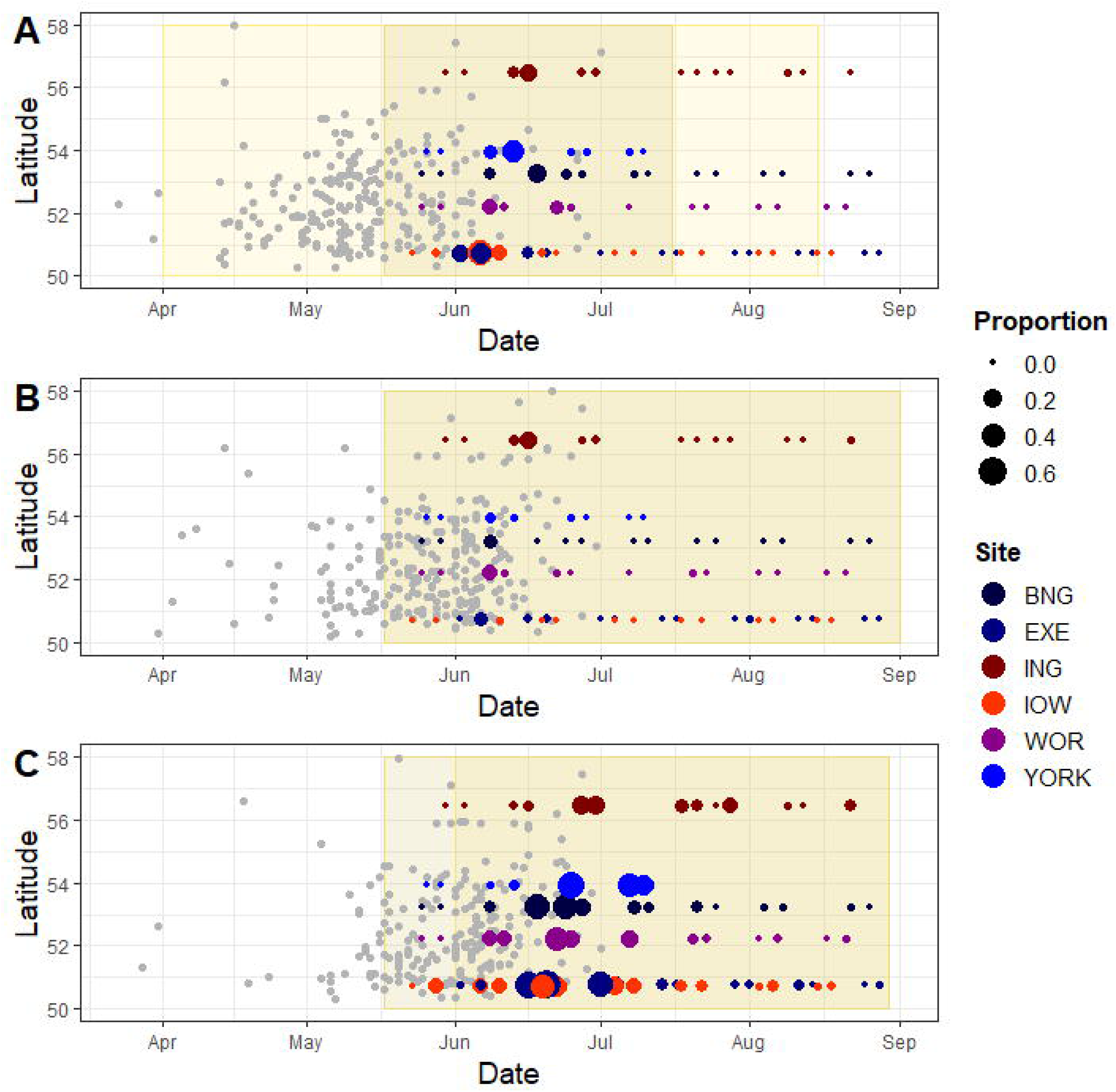
Airborne grass pollen observed 3-4 weeks after first flowering dates. Comparison of genus incidence in metabarcoding data with records of first flowering dates in 2016 from the citizen science project Nature’s Calendar (www.naturescalendar.org.uk) for (A) *Alopecurus pratensis*, (B) *Dactylis glomerata* and (C) *Holcus lanatus*. Each grey point represents the earliest time of flower heading as observed by a participant in the project. Coloured points represent metabarcoding samples, with the size of the point representing the proportion of total reads assigned to the relevant genus. Yellow shaded areas represent the expected flowering period as described in [37], with darker shades showing the ‘main’ flowering period.

Enabled by contemporary molecular biodiversity analytical approaches and mature, curated DNA barcoding databases, here we provide a comprehensive taxonomic overview of airborne grass pollen distribution, throughout an entire allergy season and across large geographic scales. The grass pollen season is defined by discrete temporal windows of different grass species, with some grass species displaying geographical variation. Temporal pollen distributions in metabarcoding data follow observed flowering times. The data provide an important step towards developing species-level grass pollen forecasting. Additionally, the research presented here leads the way for future studies facilitating understanding of the relationships between grass pollen and disease, which have significant global public health relevance and socioeconomic importance.

## Methods

### Sampling and Experimental Design

We collected aerial samples from six sites across the UK (S3 Table; S1 Fig) using Burkard Automatic Multi-Vial Cyclone Samplers (Burkard Manufacturing Co. Ltd. Rickmansworth, UK). The volumetric aerial sampler uses a turbine to draw in air (16.5 litres/min) and aerial particles are collected, using mini-cyclone technology into 1.5 ml sterile microcentrifuge tubes located on a carousel (S5 Fig). Each sampling unit was mounted alongside a seven-day volumetric trap (Burkard Manufacturing Co. Ltd. Rickmansworth, UK) belonging to the Met Office UK Pollen Monitoring Network, which provided daily pollen count data. In the seven-day volumetric trap, aerial particles are collected onto an adhesive coated tape supported on a clockwork-driven drum. The tape is cut into 24 h sections and pollen are identified and counted under a microscope [7]. Bangor was the only sampling site which was not part of the pollen monitoring network, but we deployed the same methodology at the Bangor site.

Sampling began in late May 2016 (S4 Table) and during alternate weeks, aerial samples were collected for seven days for a total of seven weeks between 25^th^ May and 28^th^ August. Exact sampling dates varied slightly between sites (S4 Table) and a total of 279 aerial samples were collected.

### DNA Extraction, PCR and Sequencing

From the 279 daily aerial samples, 231 were selected for downstream molecular analysis, as described below. Within each sampling week, two series of three consecutive days were pooled. Pooled samples were selected based on grass pollen counts obtained by microscopy. The final, unselected, day was not used in downstream molecular analysis. In total, seventy-seven pools of DNA were created. In one instance, three consecutive days of pollen samples were unavailable (Invergowrie, week 2, pool 2) due to trap errors. For this sample, the next sampling day was selected for pooling (S4 Table). DNA was extracted from daily samples using a DNeasy Plant Mini kits (Qiagen, Valencia, CA, USA), with some modifications to the standard protocol as described by [27]. DNA from daily samples was pooled and eluted into 60 µl of elution buffer at the binding stage of the DNeasy Plant Mini kit.

Illumina MiSeq paired end indexed amplicon libraries were prepared following a two-step protocol as recommended by the manufacturer [28]. Two marker genes were amplified with universal primer pairs *rbcL*af and *rbcL*r506 [19, 29], and ITS2 and ITS3 [14] (S6 Table). A 5’ universal tail was added to the forward and reverse primers and a 6N sequence was added between the forward universal tail and the template-specific primer, which is known to improve clustering and cluster detection on MiSeq sequencing platforms [30] (Integrated DNA Technologies, Coralville, USA). Round 1 PCR was carried out in a final volume of 25 µL, including forward and reverse primers (0.2 µM), 1X Q5 HS High-Fidelity Master Mix (New England Biolabs) and 1 µL of template DNA. Thermal cycling conditions were an initial denaturation step at 98 °C for 30s; 35 cycles of 98 °C for 10s, 50 °C for 30s, 72 °C for 30s; and a final annealing step of 72 °C for 5 minutes. Products from the first PCR were purified using Agencourt AMPure XP beads (Beckman Coulter) with a 1:0.6 ratio of product to AMPure XP beads.

The second round PCR added the unique identical i5 and i7 indexes and the P5 and P7 Illumina adaptors, along with universal tails complementary to the universal tails used in round 1 PCR (S4 Table, S5 Table) (Ultramer, by IDT, Integrated DNA Technologies). Round 2 PCR was carried out in a final volume of 25 µL, including forward and reverse index primers (0.2 µM), 1X Q5 HS High-Fidelity Master Mix (New England Biolabs) and 5 µL of purified PCR product. Thermal cycling conditions were: 98 °C for 3 min; 98 °C for 30 s, 55 °C for 30 s, 72 °C for 30 s (10 cycles); 72 °C for 5 min, 4 °C for 10 min. Both PCRs were run in triplicate. The same set of unique indices were added to the triplicates which were then pooled following visual inspection on an agarose gel (1.5%) to ensure that indices were added successfully. Pooled metabarcoding libraries were cleaned a second time using Agencourt AMPure magnetic bead purification, run on an agarose gel (1.5%) and quantified using the Qubit high sensitivity kit (Thermo Fisher Scientific, Massachusetts, USA). Positive and negative controls were amplified in triplicate with both primer pairs and sequenced alongside airborne plant community DNA samples using the MiSeq. Sequence data, including metadata, are available at the Sequence Read Archive (SRA) using the project accession number SUB4136142.

### Bioinformatic Analysis

Initial sequence processing was carried out following a modified version of the workflow described by de Vere *et al.* [27]. Briefly, raw sequences were trimmed using Trimmomatic v0.33 (*42*) to remove short reads (<200bp), adaptors and low quality regions. Reads were merged using FLASH v 1.2.11 [27, 31], and merged reads shorter than 450bp were excluded. Identical reads were merged using fastx-toolkit (v0.0.14), and reads were split into ITS2 and *rbcL* based on primer sequences.

To prevent spurious BLAST hits, custom reference databases containing *rbcL* and ITS2 sequences from UK plant species were generated. While all native species of the UK have been DNA barcoded [19], a list of all species found in the UK was generated in order to gain coverage of non-native species. A list of UK plant species was generated by combining lists of native and alien species [32] with a list of cultivated plants obtained from Botanic Gardens Conservation International (BGCI) which represented horticultural species. All available *rbcL* and ITS2 records were downloaded from NCBI Genbank, and sequences belonging to UK species were extracted using the script ‘creatingselectedfastadatabase.py’, archived on GitHub.

Metabarcoding data was searched against the relevant sequence database using blastn [33], via the script ‘blast_with_ncbi.py’. The top twenty blast hits were tabulated (‘blast_summary.py’), then manually filtered to limit results to species currently present in Great Britain. Reads occurring fewer than four times were excluded from further analysis. All scripts used are archived on GitHub: https://doi.org/10.5281/zenodo.1305767.

## Statistical Analysis

To understand how the grass pollen composition changed with space and time, the effect of time (measured as the number of days after the first sampling date), latitude and longitude of sampling location were included in a two-tailed generalized linear model using the ‘manyglm’ function in the package ‘mvabund’ [34]. The proportion of sequences was set as the response variable; proportion data was used as this has been shown to be an effective way of controlling for differences in read numbers [35]. The effect of time, latitude, longitude, month (coded as a factor), and the interaction between time and latitude were included as explanatory variables in the models. In addition to these explanatory variables,the interaction between time and longitude was included in a model to analyse the ITS2 data (S6 Table).

The data best fit a negative binomial distribution, most likely due to the large number of zeros (zeros indicate that a grass genus is absent from a sampling location), resulting in a strong mean-variance relationship in the data (S6 Fig). The proportion of sequences was scaled by 1000 and values were converted to integers so that a generalized linear model with a negative binomial distribution could be used. Model selection was based by Akaike Information Criterion (AIC) (S6 Table) and visual inspection of the residuals against predicted values from the models (S7 Fig).

In order to compare the metabarcoding data with flowering time data, we used phenological records of first flowering collected in 2016 by citizen scientists from the UK’s Nature’s Calendar (www.naturescalendar.org.uk). First flowering time was compared to genus-level ITS2 metabarcoding data for three species: *Alopecurus pratensis, Dactylis glomerata* and *Holcus lanatus*. As grass pollen could only be reliably identified to genus level in the metabarcoding data, the taxa compared may not have been exactly equivalent since both *Alopecurus* and *Holcus* contain other widespread species within the UK. However, *Alopecurus pratensis* and *Holcus lanatus* are the most abundant species within their respective genera. The comparison was only carried out for ITS2 data because two of the three genera were not identified by the *rbcL* marker.

NMDS ordination was carried out using package ‘VEGAN’ in R [36], based on the proportion of total high-quality reads contributed by each grass genus, using Bray-Curtis dissimilarity (S2 Fig). Ordination is used to reduce multivariate datasets (e.g. abundances of many species) into fewer variables that reflect overall similarities between samples. A linear model was carried out using the ‘lm’ function within the ‘stats’ package in R, in order to investigate the relationship between the number of reads obtained for each genus using the rbcL and ITS2 marker.

## Acknowledgements

We thank John Kenny, Pia Koldkjær, Richard Gregory, and Anita Lucaci of the Liverpool Centre for Genomic Research for sequencing support. We acknowledge the computational services & support of the Supercomputing Wales project, which is part-funded by the European Regional Development Fund (ERDF) via Welsh Government. We thank the Botanic Gardens Conservation International (BGCI) for access to the list of plant collections in the National Gardens in the UK and Ireland. We thank the Met Office network for providing additional observational grass pollen count data and Jonathan Winn, UK Met Office for ArcGIS assistance on S1 Fig. We are grateful to the Woodland Trust and Centre for Ecology & Hydrology for supplying the UK Phenology Network data and to the citizen scientists who have contributed to the latter scheme. Final thanks to Wendy Grail and technical support staff at Bangor University.

## Author Contributions

S.C., N.dV., G.W.G., R.N.M., N.J.O., C.A.S., Y.C. and G.L.B. conceived and designed the study; B.A-G., G.L.B., G.P., A.E., R.N., S.P., K.S., and N.S. collected samples and counted pollen; G.L.B. performed laboratory work, supported by S.C.; N.dV., C.R.F., L.J. and S.C. contributed methods; C.P. and G.L.B. analysed the data and G.L.B., C.P. and S.C. produced the first draft of the manuscript. All authors contributed substantially to the final submitted manuscript.

## Funding

This work was supported by the Natural Environment Research Council (https://nerc.ukri.org/), awarded to SC (NE/N003756/1), CS (NE/N002431/1), NO (NE/N002105/1) and NdV and GG (NE/N001710/1). The funders had no role in study design, data collection and analysis, decision to publish, or preparation of the manuscript.

**Competing interests:** The authors are not aware of any competing interests.

**Data and materials availability:** All sequence data (including metadata) are available at the Sequence Read Archive (SRA) using the project accession number SUB4136142. Archived sequence data was used to generate Fig 1 to 3 (including S2-S4 and S6-S7 Figs). First flowering data used in Fig 3 was obtained from Nature’s Calendar, Woodland Trust and is available upon request. The sequence analysis pipeline is available at https://github.com/colford/nbgw-plant-illumina-pipeline.

